# Sensory representations in primary visual cortex are not sufficient for subjective imagery

**DOI:** 10.1101/2024.01.10.574972

**Authors:** Giulia Cabbai, Chris Racey, Julia Simner, Carla Dance, Jamie Ward, Sophie Forster

## Abstract

The contemporary definition of mental imagery is characterized by two aspects: a sensory representation resembling, but not resulting from, perception, and an associated subjective experience. Neuroimaging demonstrated imagery-related sensory representations in primary visual cortex (V1) that show striking parallels to perception. However, it remains unclear whether these representations always reflect subjective experience, or they can be dissociated from it. We addressed this question by comparing sensory representations and subjective imagery among visualizers and aphantasics, the latter with an impaired ability to experience imagery. Importantly, to test for the presence of sensory representations independently of the ability to generate imagery *on demand* we examined both spontaneous and voluntary imagery forms.

Using multivariate fMRI, we tested for decodable sensory representations in V1 and subjective visual imagery reports that occurred either spontaneously (during passive listening of evocative sounds) or in response to the instruction to *voluntarily* generate imagery of the sound content (always while blindfolded inside the scanner). Among aphantasics, V1 decoding of sound content was at chance during voluntary imagery, and lower than in visualizers, but it succeeded during passive listening, despite them reporting no imagery. In contrast, in visualizers, decoding accuracy in V1 was greater in voluntary than spontaneous imagery (while being positively associated with the reported vividness of both imagery types). Finally, for both conditions, decoding in precuneus was successful in visualizers but at chance for aphantasics. Together, our findings show that V1 representations can be dissociated from subjective imagery, while implicating a key role of precuneus in the latter.

---

An influential definition describes mental imagery as occurring ‘*when a representation of the type created during the initial phases of perception is present but the stimulus is not actually being perceived; such representations preserve the perceptible properties of the stimulus and ultimately give rise to the subjective experience of perception’* (Kosslyn, Thompson & Ganis, 2006). This definition identifies two main components of imagery: the sensory representation, and the accompanying subjective experience. Historically, the latter has been seen as a necessary component in imagery, with the earliest definitions and long-standing measures being subjective (Perky, 1910; Richardson, 1969; Finke, 1989; Thomas, 2008; Marks, 1973). However, following the rise of cognitivism, the focus of empirical investigation in one influential thread of mental imagery research shifted progressively towards the “representational” component of the definition (Kosslyn, Ganis & Thompson, 2001). Neuroimaging research, in particular, has increasingly provided striking evidence for common neural representation of external perception (seeing objects in the world) and mental imagery (visualizing objects in the mind’s eye; Pearson, 2019; Dijkstra, Bosch, & van Gerven, 2019). Multivariate pattern fMRI approaches revealed overlapping neural representations for specific visual stimuli that are imagined versus perceived (e.g., Cichy, Heinzle & Haynes, 2012; Albers et al., 2013), to the extent that low-level features of an imagined object can be decoded from activity patterns in primary visual cortex (V1; Naselaris et al., 2015). These findings led to the conceptualization of visual mental imagery generation as “vision in reverse” (Pearson et al., 2015; Pearson, 2019; although see Spagna et al., 2021 for an alternative model), whereby activity in high-level frontal and parietal brain areas constitute the input to visual cortex, which represents the end point where the sensory representations of imagery are generated.

As noted by Nanay (2021), the majority of neuroimaging demonstrations of ostensibly imagery-related sensory representations in early visual cortex did not directly measure the accompanying subjective experience – rather, this was simply *assumed* to occur in response to task instructions to “imagine” (e.g., Albers et al., 2013; Naselaris et al., 2015), or when performing tasks that are often *thought* to involve imagery (e.g., mental rotation, Iamshchinina et al., 2021; but see Logie et al., 2011). As such, the precise relationship between the two postulated components of mental imagery – the sensory representation *in the brain* and the accompanying subjective quasi-perceptual experience *in the mind’s eye* – remains unclear. In particular, is subjectively experienced imagery always the consequence of a sensory representation in visual cortex (as implied by Kosslyn et al., 2006), or are these two components of imagery dissociable? This question formed the focus of the present research.

The subjective experience of imagery is typically measured in terms of its “vividness”, which reflects the clarity and liveliness of the quasi-perceptual experience (Marks, 1973; McKelvie, 1995). There are substantial individual differences in visual imagery vividness among the general population (Galton, 1880; Marks, 1999), with approximately 4% of the population (Dance, Ipser & Simner, 2022) struggling to experience imagery - a condition called *aphantasia* (Zeman, Dewar, & Della Sala, 2015). Due to their markedly impaired ability to experience visual mental images (despite having no deficit in visual perception), aphantasics provide a way to disentangle the relationship between sensory representations and the subjective experience of imagery itself. Indeed, the lack of theoretical agreement on the relationship between these two key components of imagery is reflected in the conflicting views regarding the underlying cause of aphantasia (see Lorenzatti, 2023, for a recent summary of these views). The ‘vision in reverse’ framing of visual cortex representation as the end point for visual mental imagery generation might imply that its subjective experience would occur as an automatic consequence of reaching this final stage. By this account, aphantasia might result from an absence of the underlying low-level sensory representation within visual cortex – for perspectives on aphantasia in line with this view see Keogh & Pearson (2018), and Wicken et al. (2021). However, this view is somewhat challenged by findings that aphantasics show preserved performance on objective tasks that presumably involve internally-generated visual cortex representations (e.g., mental rotation, Zeman et al., 2010; Pounder et al., 2022; visual working memory, Keogh, Wicken & Pearson, 2018; contingent capture, Cabbai et al., 2023). Another perspective more compatible with these findings is that the two postulated components of imagery (in terms of sub-personal mechanisms and phenomenological imagery) may be dissociable (Phillips, 2014; Faw, 2009; see also Lorenzatti, 2023). Were this the case, aphantasics might still have intact underlying sensory representations in visual cortex while entirely lacking the accompanying subjective experience.

To date, there is some initial evidence linking self-reported imagery vividness to the degree to which V1 representations overlap between imagination and perception tasks (e.g., Cui et al., 2007; Dijkstra et al., 2017). However, as these studies examined variation among people who commonly experience at least some level of imagery, they do not speak directly to the question of whether sensory representations can be dissociated from the subjective experience of imagery itself. Furthermore, the only fMRI study comparing aphantasics with visualizers during what was assumed to be a mental imagery task (“thinking about” faces and places) did not find any differences in overall BOLD activation of visual cortex (Milton et al., 2021; although they did not test for sensory representations). There are also neuropsychological studies that speak more broadly to the role of visual cortex in imagery, with mixed results (e.g., Farah, 1988; see Spagna et al., 2021 for a recent meta-analysis). Notably, there exists one demonstration of preserved subjective imagery experience in a cortically blind patient (Bridge et al., 2014), which might initially appear compatible with a dissociation between V1 representation and subjective imagery experience. However, despite showing severely disrupted V1 BOLD response to external visual stimulation, the patient showed similar activation of V1 to controls during voluntary imagery of houses – as such it remains possible that the patient was capable of generating a V1 representation. Indeed, no neuropsychological study to our knowledge has tested for both multivariate V1 representations and subjective imagery experience. Hence, the dissociability of subjective imagery experience from visual cortex representations has not yet been directly tested empirically.

A dissociation would have important implications for the field of mental imagery research, in terms of its definition as well as the way in which it has been (and should be) investigated. In particular, it would highlight that ‘imagery’ studies measuring visual cortex representations but not subjective experience may in fact reflect different underlying constructs than ‘imagery’ studies using subjective measures alone. Furthermore, a question would then remain as to what is driving the difference between people who report to experience subjective imagery and those who cannot experience any subjective imagery. Here we considered that, in the event of a dissociation with visual cortex, subjective experience differences might instead be driven by a higher-level region implicated in the subjective experience of internally-generated representations. To test this, we additionally investigated the precuneus, a central node in the default mode network (Utevsky et al., 2014). We selected precuneus as our candidate region given its well established links to the subjective experience of a variety of phenomena linked to internally-generated representations, e.g., mind wandering (Fox et al., 2015), dreams (Siclari et al., 2017), memory (Mazzoni et al., 2019). Moreover, this region has been also linked to metacognitive ability in episodic memory (Ye et al., 2018) and recovery of consciousness in patients (Wu et al., 2022). In addition, increased activation of precuneus has been found during voluntary imagery relative to perception or baseline (Fletcher, 1995; Fulford et al., 2018; Milton et al., 2021).

While the previous literature on mental imagery has almost exclusively relied on voluntary imagery paradigms, looking solely at voluntary imagery would not adequately address our research question, for several reasons. First, voluntary imagery paradigms involve an additional requirement above and beyond simply experiencing imagery: namely, the ability to generate imagery on demand. As such, any absence of sensory representations in visual cortex could potentially reflect a deficit specific to the *manipulation and control* of mental imagery. In reality, mental images in daily life also often occur seemingly spontaneously – for example, when the sound of a friend’s voice on the phone spontaneously elicits the image of their face. While it is well known that voluntary visual imagery is impaired in aphantasics, some have questioned whether spontaneous imagery might be spared (Zeman et al., 2015, 2020; Blomkvist, 2023; Cavedon-Taylor, 2022; Palermo et al., 2022). Somewhat in line with this is evidence that some aphantasics report experiencing visual imagery during dreams (Zeman et al., 2015, 2020). However, there is as yet no direct empirical evidence as to whether they also experience spontaneous imagery (or spontaneous internally-generated V1 representations) during wakefulness^1^. We therefore sought to provide the first direct test for this possibility. Importantly, this allowed us to distinguish between a deficit in the ability to form any imagery at all (in terms of either sensory representation or subjective experience), versus the ability to do so on demand.

Another reason for examining automatically-triggered ‘spontaneous’ imagery is that directly recruiting aphantasics in the light of their inability to generate imagery at will and subsequently asking them to generate imagery increases risks of demand characteristics (Cabbai et al., 2023), due to differences in expectations and motivation. Relatedly, it appears possible that the act of attempting to generate imagery on demand might itself produce a different pattern of activity among aphantasics versus visualizers (e.g., due to frustration or the adoption of alternative cognitive strategies). This itself might potentially disrupt the creation of sensory representations in V1. In other words, asking aphantasic participants to do something ‘impossible’ might paradoxically impede their capacity to form sensory representations in situations where these might otherwise arise spontaneously. As such, a full test for dissociation requires a paradigm that would also allow to measure any automatically-triggered imagery.

Not much is known about the neural correlates of spontaneous visual imagery, despite its presence in everyday life (Radomsky et al., 2014) and central role in many psychiatric disorders (Holmes & Mathews, 2005, 2010). Indeed, investigating spontaneous imagery is particularly challenging, since it is difficult to manipulate and control its content in an experimental setting. However, a promising potential approach is suggested by the work of Vetter, Smith & Muckli (2014). In their paradigm, participants were asked to listen to a series of naturalistic sounds (e.g., people chatting, birds singing, etc.) while blindfolded in the scanner. The authors found that passive listening of sounds induces a reliable sensory representation specific to the sound content in early visual cortex (e.g., the V1 pattern of activity when listening to birds singing could reliably be distinguished from that elicited by listening to people chatting). As no measure of subjectively experienced imagery was included in this prior work (since imagery was not their focus), it is unclear whether or not these spontaneous early visual cortex representations were accompanied by spontaneous visual imagery experience of the sound content (e.g., visual imagery of birds from the sound of birds singing). Nevertheless, this prior research suggests that evocative naturalistic sounds are effective means to trigger spontaneous early visual cortex representations. In the present study we therefore employed evocative sounds as a tool to investigate the relationship between sensory representations and subjective experience, in relation to both spontaneous (elicited during passive listening of sounds) and voluntary visual imagery (elicited on demand) among visualizers and aphantasics. Notably, an exploratory searchlight analysis in Vetter et al. (2014) revealed that decodable representations of sound (as observed in V1) were also present in precuneus. The authors originally interpreted the role of the precuneus as a hub mediating between the auditory and visual representations of sounds. However, in light of the aforementioned evidence, an alternative interpretation could be that precuneus mediates the translation of internally-generated V1 sensory representations into the subjective experience of imagery. By directly measuring the subjective experience of imagery (which Vetter et al. did not measure), our study allows us to address this possibility.

Across three pilot studies, we first developed a set of sounds that would be effective in inducing the subjective experience of spontaneous imagery (among those who can experience it). We then recruited 26 visualizers and 24 aphantasics and asked our participants to listen, blindfolded, to the series of evocative sounds (e.g., sounds of cat meowing, dog barking, traffic noises, etc.) in two different conditions, while fMRI data was acquired. In the first condition, designed to elicit spontaneous imagery (at least, among those capable of this), participants were just asked to listen carefully to the sounds without receiving any other instruction (passive listening task, as in Vetter et al., 2014). Importantly, during this task participants were completely naive to the fact that it was designed to trigger visual imagery (as this would inevitably impact the “spontaneity” of the imagery being generated). Instead, at the end of this passive listening task, participants received a surprise question asking them whether they spontaneously experienced visual mental imagery in response to the sounds. Subsequently, participants completed the voluntary imagery task, which was identical to the first task but with the addition of explicit instruction to generate voluntary visual imagery of the sounds content. We also separately collected ratings of voluntary mental imagery vividness. Using this method, we were able to assess the relationship between sensory representations in V1 and the subjective experience of imagery that occurred both spontaneously (i.e., during passive listening) and voluntarily, among both aphantasics and visualizers. The strength of sensory representations was indexed through the accuracy with which a linear support vector machine algorithm could classify (or ‘decode’) the contents of different sounds, through their associated specific pattern of V1 activity.

In particular, we predicted that, if the two components of imagery are always associated, aphantasics’ decoding accuracy in V1 should be at chance level (reflecting a lack of decodable low-level sensory representations), and lower than in visualizers (where we expected above chance level decoding). And it should be this way during both passive listening and voluntary imagery. A modified version of this view could be that, if imagery deficits in aphantasia are restricted to voluntary imagery only (Zeman et al., 2015), then aphantasics might report spontaneous (but not voluntary) imagery, and produce decodable V1 patterns in the former condition only. On the other hand, if the two components can be dissociated, we expected aphantasics to show decodable sensory representations in V1, but without reporting any associated imagery experience, in at least one of the two conditions. Were this the case, we reasoned that aphantasics might instead lack representations in precuneus.

## Results

### Group differences in both spontaneous and voluntary forms of subjective imagery experience

First, we assessed whether vividness ratings for spontaneous and voluntary imagery elicited by sounds varied between aphantasics and visualizers. We found a main effect of group, *F*(1, 48) = 167.07, *p* < .001, 𝜂²_p_ = 0.78, but not imagery type (spontaneous, voluntary), *F*(1, 48) = 2.68, *p* = .108, 𝜂²_p_ = 0.05, or interaction, *F*(1, 48) = 0.83, *p* = .37, 𝜂²_p_ = 0.02. Figure 1B shows that aphantasics’ mean vividness ratings of voluntary (*M* = 1.07, *SD* = 0.2) and spontaneous (*M* = 1, *SD* = 0) imagery were lower than visualizers (voluntary rating *M* = 3.01, *SD* = 0.5, spontaneous rating *M* = 2.77, *SD* = 1.07). Our sounds were effective at inducing spontaneous subjective imagery in the majority (22/26) of visualizer participants, all of whom *also* reported imagery during the voluntary condition. In contrast, all 24 aphantasics rated spontaneous imagery vividness as equal to 1 (“no image at all”), while only four out of 24 had a mean vividness of voluntary imagery different from 1 (but still within the typical aphantasia vividness range, i.e., rated as somewhere between entirely absent or vague/dim/fleeting, Zeman et al., 2015, 2020; Dance et al., 2021, 2022).

**Figure 1.**
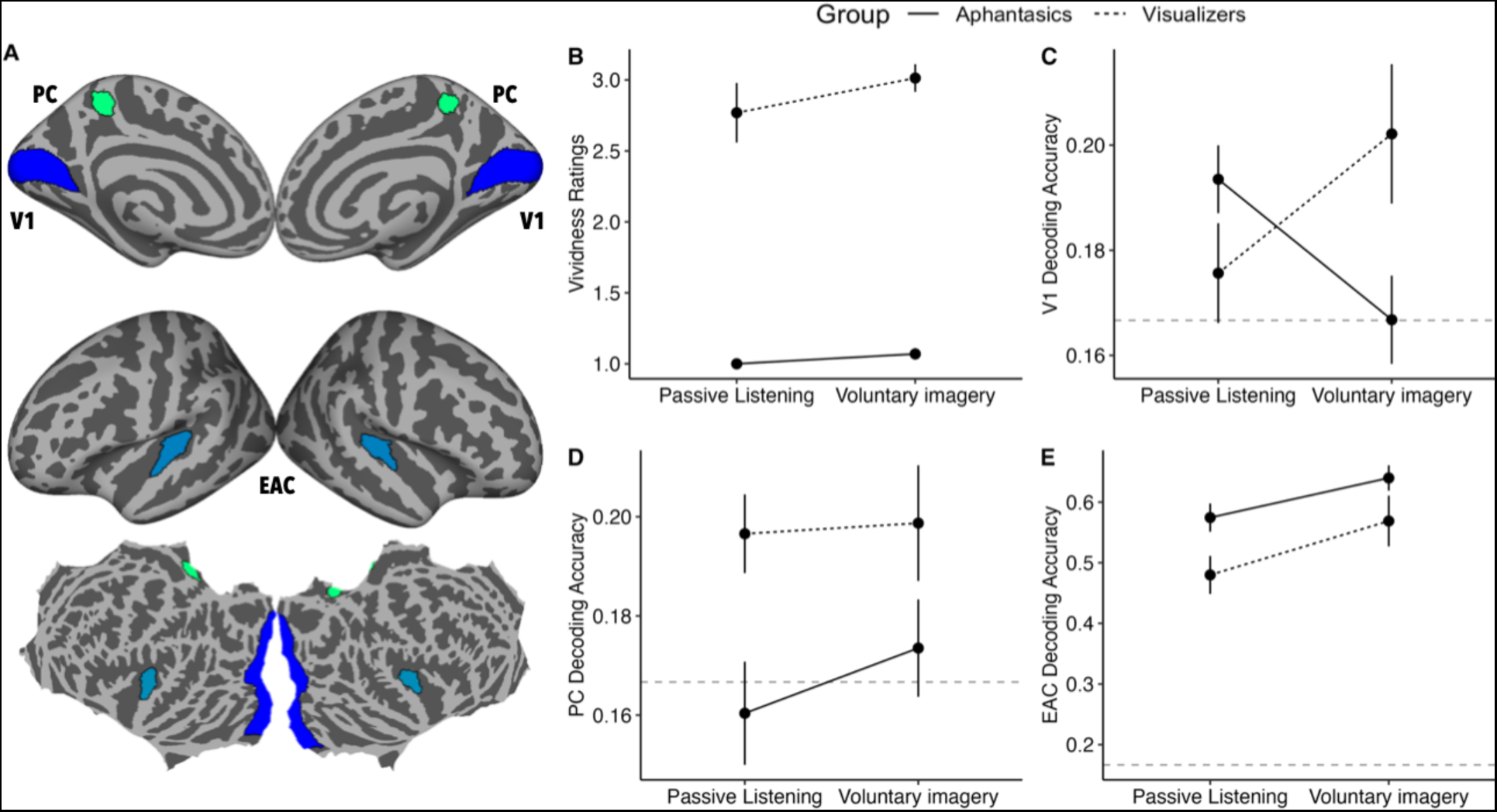
***A.*** Medial, lateral and flat view of our regions of interest (ROIs): primary visual cortex (V1), precuneus (PC), early auditory cortex (EAC), indicated by three different colors. We mainly focused on V1 and PC for addressing our research questions, whereas EAC was used as control region. ***B.*** Vividness ratings about the sounds as a function of group (aphantasics, visualizers) and imagery type (passive listening task, reflecting spontaneous imagery, and voluntary imagery task). Spontaneous imagery ratings consisted of a single rating provided at the beginning of the vividness ratings task, whereas voluntary imagery ratings were provided for each sound, and averaged for each participant. ***C, D, E.*** Decoding accuracy of sounds categories in V1 (*C*) PC (*D*) EAC (*E*) as a function of task condition (passive listening, reflecting spontaneous imagery, and voluntary imagery task) and group (aphantasics, visualizers). Dashed horizontal line indicates chance level (⅙ sounds categories). Error bars show SEM.

### V1: opposite patterns for spontaneous and voluntary representations in aphantasics vs. visualizers and the link with subjective ratings of vividness

We applied multivariate pattern analysis (MVPA) to fMRI data to test for the presence of stimulus-specific sensory representations in V1 among aphantasics and visualizers. Specifically, we tested for these representations in relation to the five functional runs in the spontaneous imagery condition (passive listening task) and the five functional runs in the voluntary imagery condition (i.e., elicited by the instruction to form a mental image). During each run, participants listened to 18 natural sounds (each belonging to one of 6 categories, e.g., dog barking, seagull noises, etc.) interleaved by silence. We first extracted betas for each voxel in V1, representing the estimates of the trial-wise BOLD response amplitudes to each stimulus (4 seconds of sound presentation) relative to the BOLD signal observed during the absence of a stimulus (silence). To decode activity patterns reflecting sound content in V1, we fed these estimates to a linear support vector machine classification algorithm that was trained on all but one run to discriminate between the 6 sound categories and tested on the remaining run (‘leave-one-run-out’ cross-validation).

We then compared decoding accuracy of sound categories in V1 between our two groups of participants and tasks. There was no significant main effect of group, *F*(1, 48) = 0.62, *p* = .44, 𝜂²_p_ = 0.01, or task, *F*(1, 48) = 0.00, *p* = .99, however, the interaction between the two was significant, *F*(1, 48) = 9.85, *p* = .003, 𝜂²_p_ = 0.17. This reflected that the difference between aphantasics and visualizers emerged only for the voluntary imagery task, *t*(89.7) = −2.53, *p* = .01, with aphantasics’ decoding accuracy (*M* = 0.17, *SD* = 0.04) being lower than visualizers (*M* = 0.20, *SD* = 0.07), see Figure 1C. On the other hand, no group difference emerged for the passive listening task, *t*(89.7) = 1.27, *SE* = 0.01, *p* = .20. As shown in Figure 1B, decoding accuracy within aphantasics was lower for the voluntary imagery task (*M* = 0.17, *SD* = 0.04) than for passive listening (*M* = 0.19, *SD* = 0.03), *t*(48) = 2.19, *p* = .034, whereas in visualizers, decoding accuracy was higher for the voluntary imagery task (*M* = 0.20, *SD* = 0.07) than for passive listening (*M* = 0.18, *SD* = 0.05), *t*(48) = −2.25, *p* = .029.

Indeed, among visualizers, decoding accuracy in V1 was significantly above chance for the voluntary imagery task, *p* < .001, but did not reach the threshold of a significant difference from chance level during the passive listening task, *p* = .147. We note that, in this group, not all participants reported experiencing spontaneous subjective imagery during passive listening. When removing the 4 visualizers who reported experiencing no imagery, classification accuracy was above chance (*p* = .044). As shown in Figure 2A, in visualizers, decoding accuracy during this task (passive listening) was positively related to vividness ratings of spontaneous imagery, *r*(24) = .46, *p* = .017, suggesting that the strength of sensory representations in V1 was linked (at least in part) to spontaneous subjective imagery experience. Consistently with this, in visualizers we also found a positive correlation between decoding accuracy in V1 during the voluntary imagery task and voluntary imagery vividness ratings, *r*(24) = .48, *p* = .012 (Figure 2B).

**Figure 2.**
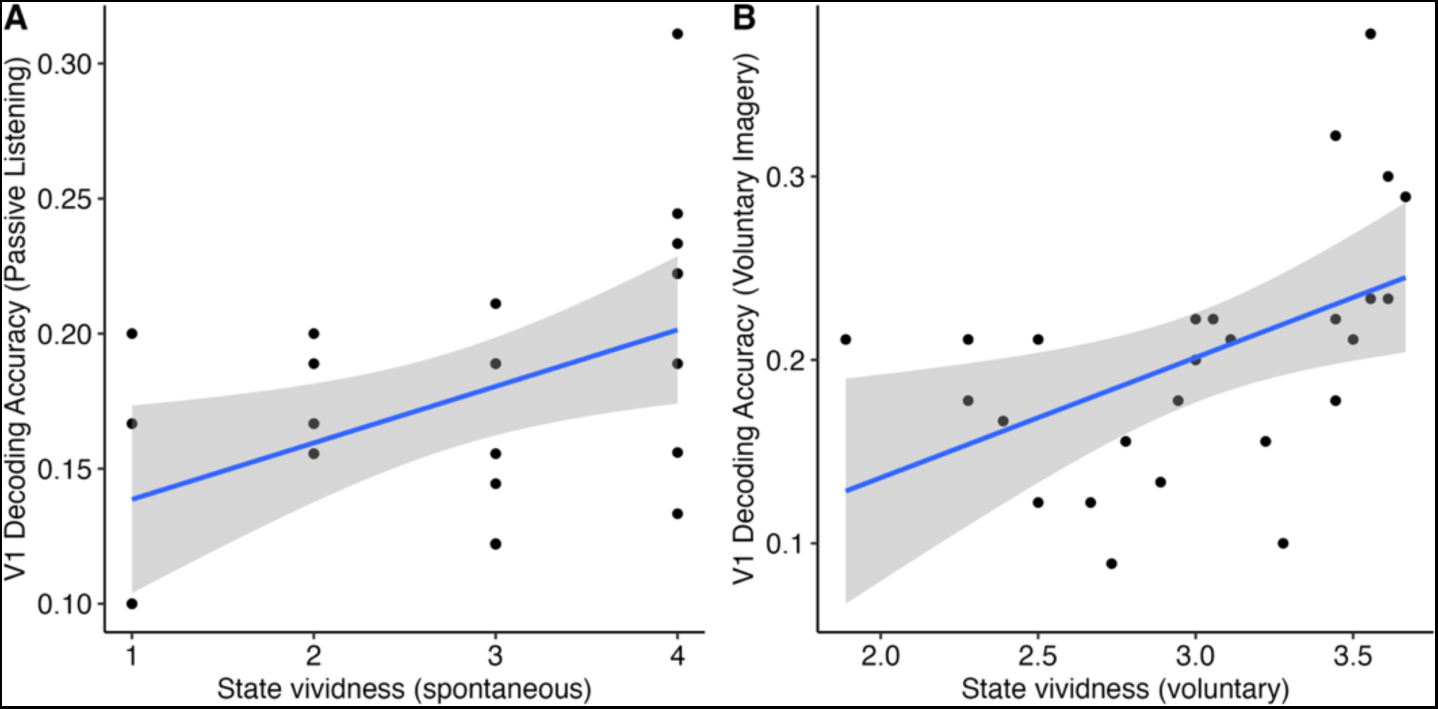
Decoding accuracy of natural sounds in V1 in visualizers was positively correlated with ***A.*** subjectively experienced mental imagery vividness during both passive listening (reflecting spontaneous imagery, *r* = .46) and ***B.*** while generating voluntary imagery related to the sounds content (*r* = .48). Shaded interval shows 95% CI.

In contrast to the visualizer group, aphantasics’ decoding accuracy was higher than chance level for the passive listening task, *p* < .001, but not for the voluntary imagery task, *p* = .51. The significant decoding found in aphantasics during a condition designed to induce spontaneous imagery was in stark contrast with the absence of spontaneous imagery in self-report (vividness equal to 1). This key evidence demonstrates a dissociation between the sensory representation in V1 and the subjective experience of imagery. We note that the floor-level subjective ratings of aphantasic participants in both conditions did not permit the correlational analysis in this group.

### Precuneus: decoding above chance for visualizers but not for aphantasics in both tasks

The finding of decodable sensory representation in V1 in the absence of any conscious experience of imagery in aphantasics may suggest that involvement of higher-level regions may be necessary for the phenomenological experience of imagery. To test this, using the same analysis approach (MVPA), we examined activity patterns in our precuneus region of interest.

Here, we found a main effect of group, *F*(1, 48) = 7.32, *p* = .009, 𝜂²_p_ = 0.13, with aphantasics’ decoding accuracy (*M* = 0.17, *SD* = 0.05) being lower than visualizers (*M* = 0.20, *SD* = 0.05) in this region. This difference was due to decoding accuracy in visualizers being significantly above chance level (*p* < .001 in both tasks) while aphantasics were at chance level in both tasks (*p* = .22 for voluntary imagery and *p* = .74 for passive listening). We found no main effect of task, *F(*1, 48) = 0.79, *p* = .38, 𝜂²_p_ = 0.02, nor interaction between task and group, *F*(1, 48) = 0.41, *p* = .52, 𝜂²_p_ = 0.008, see Figure 1D.

### Early auditory cortex: decoding above chance in both group and tasks

While V1 and precuneus represented the main target of decoding analyses to address our research questions, we also investigated the auditory cortex as a control region, since a reliable decoding of sound categories has been reported in multiple studies by Vetter et al. (2014, 2020). As expected, we found higher than chance level decoding accuracy in both groups and tasks (all *p* < .001). In terms of comparisons between task and group, we found a significant main effect of task, *F*(1, 48) = 20.34, *p* < .001, 𝜂²_p_ = 0.30, indicating lower decoding accuracy during passive listening (*M* = 0.53, *SD* = 0.15) than during the voluntary imagery task (*M* = 0.60, *SD* = 0.17), see Figure 1E. Unexpectedly, we also found a significant main effect of group, *F*(1, 48) = 4.17, *p* = .047, 𝜂²_p_ = 0.08. Aphantasics were characterized by higher decoding accuracy of sound content in this region (*M* = 0.61, *SD* = 0.11) relative to visualizers (*M* = 0.52, *SD* = 0.19). No significant interaction between group and task was observed, *F*(1, 48) = 0.48, *p* = .49, 𝜂²_p_ = 0.009.

## Discussion

Our study presents novel evidence on the link between the two proposed components of mental imagery: the quasi-perceptual phenomenological experience, and sensory representations in primary visual cortex. Our results show for the first time that the subjective experience of visual imagery can be dissociated from primary visual cortex representations. Specifically, it was possible to decode information about stimulus content (e.g., dog barking, seagulls’ noises, etc.) in V1 for aphantasic participants, in the complete absence of any subjectively experienced imagery, during passive listening of evocative sounds. Indeed, decoding accuracy was similar between aphantasics and visualizers in this condition, which successfully elicited spontaneous subjective imagery in the latter group, but not in aphantasics (contrary to prior proposals e.g., Zeman et al., 2015). Also notable was that, among visualizers, the stimulus-specific pattern of V1 activity evoked in this condition was positively related to the reported vividness of imagery elicited by the sounds. This suggests V1 patterns somehow “feed up” into experienced imagery vividness in some way (for those capable of experiencing subjective imagery), even when these patterns are triggered automatically rather than through voluntary attempts to generate imagery. Crucially, however, the sole presence of decodable sensory representations in V1 is not *sufficient* to produce the phenomenological experience of imagery (as demonstrated by aphantasics). In fact, internally driven sensory representation in V1 in the absence of visual stimulation can exist even if there *is no subjective* imagery experience.

Our demonstration of a dissociation carries implications for both the definition and investigation of mental imagery. While the most long-standing and widely used measures of mental imagery index subjective experience (e.g., VVIQ, Marks, 1973), subjective experience is often not directly assessed in neuroimaging studies of (ostensibly) imagery-related sensory representations. Our results imply that studies that measure only one component of imagery and not the other (e.g., V1 representation in the absence of subjective reports, or vice versa) cannot necessarily be assumed to have investigated the same underlying construct. As such, it would be problematic to have the same term ‘mental imagery’ used interchangeably for these two dissociable components. Given the central role of subjective experience in long-standing definitions of imagery (Richardson, 1969; Finke, 1989), and the sole reliance of a large proportion of the imagery literature on such measures, this problem cannot be solved simply by dropping subjective experience from the definition of mental imagery, as proposed recently by Nanay (2021). Rather, our data highlight the need for a definitional distinction between subjective imagery experience and related sensory representations (Phillips, 2014), and specificity regarding which of these is the subject of investigation.

Our finding of a dissociation is at odds with a view of visual cortex as the endpoint of the process by which mental imagery experience is generated, as might be inferred from the “vision in reverse” model (Pearson, 2019). However, we note that while this model was primarily informed by studies of voluntary imagery, our observed dissociation arose only in our spontaneous imagery condition. Indeed, the pattern of results seen in our voluntary imagery condition aligns well with the predictions of the vision in reverse model, with both subjective imagery experience and decodable V1 representations being absent among aphantasics but present in visualisers. Also consistent with the vision in reverse model is our finding that top-down attempts to generate imagery (in the voluntary condition) boosted the strength of V1 sensory representations among visualisers, compared to the spontaneous imagery condition. Critically, however, our data highlight that considering only voluntary imagery does not give the full picture: the results of our spontaneous condition clarify that the lack of subjective imagery (in aphantasics) cannot always be explained by an absence of a sensory representation in V1. As such, results from the spontaneous condition rather support proposals that the components of imagery (i.e., subjective experience versus V1 representation) are not only distinct but can be dissociated (Phillips, 2014; Faw, 2009).

Our data demonstrate not only a dissociation between the presence/absence of V1 representation versus subjective imagery experience, but also a higher-level neural correlate of the latter. Indeed, in our precuneus ROI, decodable sensory representations were found in visualizers (across both conditions, spontaneous and voluntary imagery) but not in aphantasics. But what *exact* role might precuneus play, given our data? As noted earlier, Vetter et al. (2014)’s study of V1 decodable representations of sound also revealed, in an exploratory analysis, decodable precuneus representation. The explanation proposed by Vetter et al., which is less compatible with our data, is that precuneus acts merely as an audiovisual hub mediating between the auditory and visual representations of sounds. However, this explanation would have predicted equivalent findings across participant groups, which we did not see, and it is also incompatible with V1 and the EAC results. The function of the precuneus has been much debated (e.g., Cavanna et al., 2006), but as noted above, this region has been previously associated with imagery (Fletcher, 1995) and more broadly, subjective experience of internally generated representations such as mind wandering (Fox et al., 2015), dreams (Siclari et al., 2017), memory (Mazzoni et al., 2019; Ye et al., 2018) but also, general recovery of consciousness in patients (Wu et al., 2022). Our findings hence appear to extend this literature, as precuneus mapped onto the very presence (or absence) of subjective imagery experience. This result implies that, somewhat contrary to the conception of imagery as “vision in reverse” (Pearson, 2019), visual cortex representations may not be the endpoint for conscious imagery – rather, accompanying subjective experience appears to require the involvement of higher-level regions (with precuneus as a promising candidate). As such, precuneus may play an important role in facilitating translation of internally driven sensory representations into subjective experience of images.

Our findings also speak to theories of aphantasia. Our data show that aphantasia cannot be solely explained by an absence of underlying sensory representation in V1 (as predicted by Keogh & Pearson, 2018), but also do not support the view (at least for our sample) that aphantasics might exclusively lack subjective voluntary imagery (Zeman et al., 2015). Indeed, we found that aphantasics can generate V1 representations (in the absence of external visual stimulation) when triggered automatically, yet do not appear to “see” this information as a mental image (somewhat in line with the disconnection hypothesis discussed by Lorenzatti, 2023). It is also worth noting that the results concerning our voluntary imagery condition speak against “unconscious imagery” theories of aphantasia (Nanay, 2021). Indeed, in this condition, which most resembles how aphantasia is usually studied and assessed (by scores on the VVIQ specifically), we found both the sensory representation in V1 and subjective experience were absent in aphantasics, i.e., no form of imagery at all, whether conscious or unconscious, was observed. This, together with our finding of a brain-based objective correlate of subjective imagery presence (in precuneus), rules out the possibility that the absence of subjective imagery experience in our aphantasic sample is purely psychogenic (de Vito & Bartomoleo, 2016) or solely reflected by differences in reports (e.g., Schwitzgebel, 2011). Rather, our data points to a role of higher-level disruption (involving, although not necessarily limited to, precuneus) that may prevent internally-generated visual V1 representations from feeding into the subjective experience of mental imagery. Moreover, given the lack of subjective experience even in the spontaneous imagery condition, aphantasia disruption does not appear to be limited to the feedback mechanisms implicated in imagery generation (cf. Pearson, 2019; but also see Spagna et al., 2021). Rather, there may also potentially be disruption to the forward connectivity mechanisms (V1 to precuneus) necessary to have the phenomenological experience of imagery that accompanies visual cortex representations.

Either way, by demonstrating that aphantasics are able to spontaneously generate V1 representations, our findings offer an explanation for the once-puzzling preserved performance of aphantasics on objective behavioural tasks thought to tap into imagery-related processing (e.g., Zeman, 2010; Pounder et al., 2022; Cabbai et al., 2023; Liu & Bartolomeo, 2023). Interestingly, Palermo et al. (2022) found that aphantasics show unimpaired performance on the ‘colour imagery task’, which involves viewing black and white line drawings of fruits and vegetables and reporting which one has a darker colour. While this was interpreted by the authors as reflecting involuntary imagery, our findings suggest that aphantasics may be able to perform such tasks by drawing on their intact knowledge of visual properties without any accompanying subjective imagery experience.

An interesting question is why aphantasic participants were characterised by a reduction in decoding accuracy in V1 in the voluntary imagery condition compared to the spontaneous condition. One might have expected that, while aphantasics would not be able to ‘boost’ their V1 representations in the same manner as visualizers when explicitly instructed to visualize the sound content, they would still maintain the same spontaneously generated V1 representations of sounds in both conditions. As this didn’t occur, our findings raise the possibility that the very act of attempting to generate imagery interferes with aphantasics’ ability to form spontaneous sensory representations. The fact that aphantasics, like visualizers, showed increased rather than reduced auditory cortex decoding during the voluntary condition suggests they were still attending to the sounds, and did not simply disengage from the task (e.g., due to frustration at being asked to do the very thing that they had already told the research team they could not do, i.e., generate voluntary imagery). Another possibility is that aphantasics adopted an alternative cognitive strategy in response to the instruction to generate imagery (Zeman et al., 2010). Either could have disrupted pathways involved in the formation of V1 sensory representation reflecting sound content (e.g., dog) and hence their automatic triggering of V1 representations by sounds.

Regardless of the explanation for the negative impact of explicit imagery instructions on the ability of aphantasics to generate sensory representation in V1, this aspect of our findings highlights a potential limitation when studying the neural correlates of aphantasia, and more generally, of variation in imagery experience using only voluntary imagery paradigms. Indeed, if we only employed a voluntary imagery paradigm, we may have concluded that aphantasics cannot generate V1 sensory representations in the absence of corresponding sensory stimulation. The present work provides a novel methodology for investigating spontaneous imagery: visual imagery can be spontaneously evoked (in those who have it) by using naturalistic sounds tightly linked to specific pictures. Our approach also has the additional methodological advantage that it did not rely on mental images of stimuli that were presented just before the imagery period (e.g., as in retro cue paradigms), but rather on participants’ idiosyncratic and unconstrained visual images generated in response to sounds from long term memory. This aspect of our paradigm not only added ecological validity (as imagery in the real world mainly involves generating mental images from long-term visual memory) but also allowed us to isolate imagery from working memory traces, which was important for us since aphantasics appear to have a largely unimpaired visual working memory function (Keogh, Wicken & Pearson, 2021) and related representations in V1 (Weber et al., 2023).

In conclusion, for the first time, we provide evidence for differences between aphantasics and visualizers in sensory representations in V1 during a condition designed to measure voluntary, but not spontaneous imagery. Critically, however, stimulus-related information in V1 during passive listening (designed to elicit spontaneous imagery) was present in aphantasic participants, despite their reports of imagery absence. As such, our results establish a dissociation between the two key components of mental imagery: sensory representation in visual cortex in the absence of external stimulation, and the accompanying subjective experience. Specifically, our results demonstrate that the presence of perceptual representations in sensory cortices (such as V1) may feed into imagery experience, but is not sufficient for the subjective experience of imagery to arise. Instead, the presence or absence of subjective imagery experience was predicted by a higher-level region: precuneus. Finally, our work suggests that investigating mental imagery by considering only a single intentionality level (voluntary, as opposed to spontaneous) is not sufficient for reaching a comprehensive characterization of this phenomenon and its link with consciousness.

## Methods

### Participants

Sample size estimation was based on the appearance of large effect sizes in existing research comparing aphantasics and visualizers on imagery measures (Keogh & Pearson, 2018; Wicken et al., 2021; Zeman et al., 2015; Milton et al., 2021). For a two-tailed two-independent-group t-test with a large effect size (Cohen’s *d* = 1), with α = 0.05 and power = 0.90, the projected necessary sample size is 23 participants per group (GPower 3, Erdfelder et al., 2007). To allow for potential dropout or data loss (participants moving excessively during scanning or technical issues), 51 participants gave written informed consent and participated in the experiment (24 aphantasics and 27 visualizers). One visualizer participant was excluded because of issues with equipment (headphones), which caused them to not hear the sounds for the entire first task (passive listening). One aphantasic participant completed only 4 runs of both tasks (instead of 5) due to technical issues with the scanner.

Our final sample was composed of a group of 24 participants with aphantasia (*M*_age_ = 25.7, *SD*_age_ = 5.6, 16 females, 8 males), defined as having a mean VVIQ score equal to or lower than 2 (corresponding to a total VVIQ score of 32, Dance, Ward & Simner, 2021). The other group was constituted of 26 visualizer participants (*M*_age_ = 23, *SD*_age_ = 4.86, VVIQ mean score > 2, 18 females, 8 males). The mean VVIQ score of the aphantasic group (*M* = 1.02, *SD* = 0.09) was lower than that of the visualizer group (*M* = 3.98, *SD* = 0.55), *t*(26.51) = −27.11, *p* < .001. We note that 91% of our aphantasics were extreme aphantasics (reporting a complete absence of visual imagery). We also collected participants’ scores on the Spontaneous Use of Imagery Scale (Reisberg, Pearson, & Kosslyn, 2003), as an additional trait measure of mental imagery use in daily life (more focused imagery frequency than the VVIQ). In this measure too, aphantasics (*M* = 15.8, *SD* = 3.35) scored lower than the visualizer group (*M* = 41.3, *SD* = 4.77), *t(*44.92) = −22.04, *p* < .001.

Aphantasics were recruited via email invitation from the University of Sussex’s Imagery Lab - Aphantasia Cohort database as well as from the Aphantasia Network database. These databases were generated by recruiting aphantasics through multiple sources – those who self-referred to the Imagery Lab or Aphantasia Network to volunteer for research and those who were recruited via advertising on online forums and social media (e.g., aphantasia Facebook groups, Reddit). Visualizer participants were recruited from an existing database generated by prescreening undergraduate students at the University of Sussex on the VVIQ, by word of mouth, and via the University of Sussex Participant Recruitment database (SONA participant pool).

All participants were right-handed and reported to have normal or corrected to normal vision, normal hearing and did not have MRI contraindications. All participants received monetary compensation of £15 for their time. The study was approved by the Research Governance Ethics Committee (RGEC) at Brighton and Sussex Medical School (ER/GC337/4).

### Materials

#### Questionnaire measures

##### The Vividness of Visual Imagery Questionnaire (VVIQ; Marks, 1973)

Before scanning, all participants completed the VVIQ to assess their self-reported ability to generate visual imagery. This questionnaire is typically used to identify people with aphantasia (e.g., Milton et al., 2021; Dance et al., 2022). Participants are asked to generate visual mental images for 16 different scenarios and rate the vividness of their visual imagery on a scale from 1 (“No image at all, you only ‘know’ that you are thinking of the object”) to 5 (“Perfectly clear and as vivid as normal vision”). The questionnaire was scored by calculating the mean score of the 16 scenarios, resulting in a score ranging from 1 to 5 (and equivalent total score ranging from 16 to 80), with higher scores indicating more vivid self-reported visual imagery.

##### Spontaneous Use of Imagery Scale (SUIS; Reisberg, Pearson, & Kosslyn, 2003)

This questionnaire was also completed prior to scanning. The SUIS is a 12-item scale that measures the self-reported tendency to use mental imagery in daily life (frequency and likelihood). People are asked to indicate on a 5-point Likert scale how often a particular statement is appropriate for them, from 1 (“Never appropriate”) to 5 (“Always completely appropriate”). Each statement concerns a daily-life situation in which images are involved or may be helpful. A total score was calculated by summing responses to the 12 statements, ranging from 12 to 60, with a higher total score indicating more use of visual imagery in daily life. We included exploratory analyses related to this trait and the VVIQ in Supplementary Material.

#### Stimuli

The stimuli used in this experiment consisted of a set of short extracts (4 s) of 18 natural sounds. Specifically, each sound belonged to one out of six categories: a dog barking, a cat meowing, a crowd of seagulls, a fireplace, helicopter blades, and, finally, traffic noise. Each sound category had three different sound exemplars (e.g., the “dog” category had three different auditory stimuli representing a different dog barking). We decided to have three exemplars per sound to avoid having too many repetitions of the same auditory stimulus, which could induce participant fatigue and ultimately negatively impact on imagery being generated, or even induce adaptation (repetition suppression) and alter brain response (Grill-Spector et al., 2006). Additionally, we expected that within-category exemplars would induce very similar and unambiguous visual representations, which could nevertheless be distinguished from the other sound category exemplars in V1.

The selection of sounds was based on results from three online pilots in which we collected imageability ratings and how recognizable the sounds were to participants (N = 176 across pilots, see Supplementary Material). Stimuli were initially selected from a pre-existing database (IADS-E, Yang et al., 2018; IADS-2, Bradley & Lang, 2007a) and online sources (e.g., https://freesound.org/). All sounds’ energy (RMS) levels were equalized and presented in mono.

### Design and procedure

The main fMRI scanning session occurred at the Clinical Imaging Science Center (CISC) on the University of Sussex Campus. The experiment was programmed and presented using MATLAB R2019a (Mathworks, Inc.) with Objective Psychophysics Toolbox (Hartmann & Weisz, 2020, available at https://gitlab.com/thht/o_ptb) and Psychtoolbox (Brainard, 1997; Kleiner et al., 2007). Before entering the scanner, participants were instructed to wear a blindfold (to remove visual stimulation during some parts of the scanning session) and close their eyes. The room lights were switched off. MRI-safe in-ear headphones were used to deliver sound stimuli, and over-ear protectors were placed on them to block out scanner noise. Prior to commencing the first task, to test whether participants could clearly hear the sounds on top of the scanner noise and adjust the volume to a comfortable level, a single example sound was played (which was not part of the stimuli used for the main task) while the scanner was functioning.

The experimental session was divided into three parts (see Figure 3): functional scanning for passive listening of sounds, structural scanning that coincided with a vividness rating task (see below), and finally, functional scanning with an instruction to attempt the generation of voluntary imagery of sounds. For the first part, participants were just instructed to listen carefully to the sounds. Importantly, no reference to imagery was made at this stage. They underwent five functional runs of the passive listening task, each lasting for 3 minutes and 50 seconds (144 volumes per run). On every run, subjects listened to all the 18 auditory stimuli (4 s each) presented in a pseudo-randomized order (without repetitions), separated by an inter-stimulus interval (ISI) of 6 s, while a blank period (10 s of no sound) was presented every six sounds. The blank period was used to reduce participants’ fatigue between auditory stimuli presentations and ensure robust estimation. Each sound exemplar (e.g., three different dogs barking) was repeated five times, for a total of 15 repetitions per sound category.

**Figure 3.**
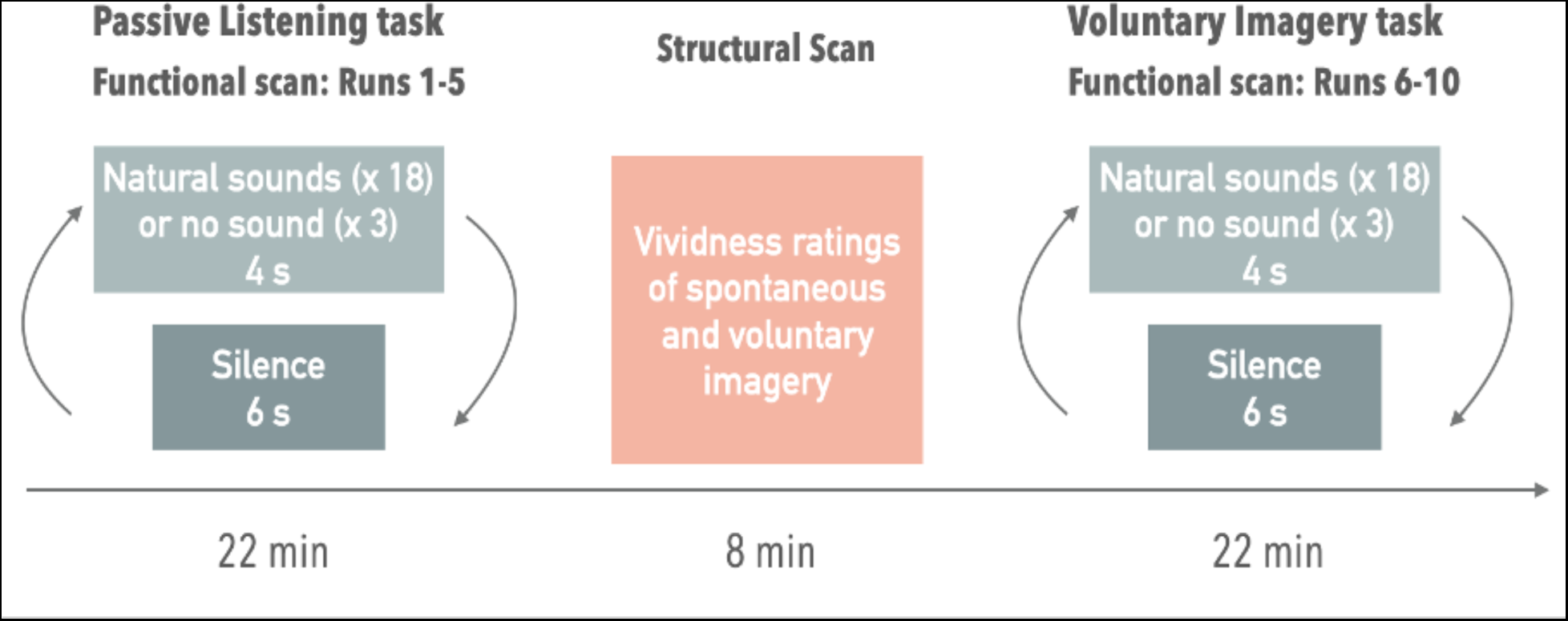
Experimental Design. On every trial, participants listened to one of 18 different natural sounds (3 exemplars for each of the 6 categories: dog, cat, traffic, helicopter, fire, seagulls), or silence (with exception of scanner noise). Each trial was separated by 6 seconds of silence (scanner noise). Completion of the vividness rating task took place during the 8 minutes structural scan.

At the end of this part, participants temporarily removed the blindfold to complete the vividness rating task administered during structural scanning. The task questions and ratings were back-projected on a screen and viewed through an angled mirror on the head coil. At the start of the task, participants were asked to answer the following surprise question “*Did you experience spontaneous visual imagery (pictures in your mind) while listening to the sounds? Please rate now the vividness of your spontaneous visual imagery from 1 (no image at all, rightmost button) to 4 (very vivid, leftmost button)*”. They were then asked to listen to all the auditory stimuli again, but this time, they were given the following instruction “*Now, please actively try to create visual imagery of what the sounds represent while listening to them. At the end of each sound, please respond to the instructions on screen which will ask you to rate the vividness of your voluntary visual imagery*”. In this way, we collected both spontaneous and voluntary imagery vividness ratings. Participants provided their ratings by pressing a button on the 4-button box. Before commencing the vividness rating task, participants were told by the experimenter that there was no right or wrong answer and that they only needed to respond as honestly as possible based on their own experiences. The vividness rating task was very short (around 3 to 4 minutes); hence, participants had time to rest for the remaining five minutes of the structural scanning.

Once the structural scanning ended, participants wore the blindfold again and underwent five additional functional runs, which coincided with the voluntary imagery task. The structure of the five runs was identical to the first part, except that, in this case, participants received explicit instruction to engage in voluntary visual imagery of the sounds while listening to them, until the end. The total scanning time for each participant was approximately 1 hour.

#### MRI data acquisition

Blood oxygen level-dependent signals were acquired using a 3T Siemens Prisma scanner with a 64-channel whole-brain coil (Siemens Medical, Erlangen, Germany). For each participant, 144 functional images per run were acquired with a gradient echo type echo planar imaging (EPI) sequence with a multiband acceleration factor of 3 with the following parameters [echo time (TE) = 28 ms; repetition time (TR) = 1520 ms; flip angle = 52 degrees; slice thickness = 2.0 isotropic mm; slices = 72; FOV = 208x180mm;]. SpinEcho Field maps with reversed phase-encode blips in both Anterior to Posterior and Posterior to Anterior were acquired with the same parameters of the functional images. Between passive listening and voluntary imagery task, a high-resolution structural scan was acquired, this was a Multiecho T1w MPRAGE (0.8mm resolution; 8 mins 22 seconds; multi-echo TE = 1.8/3.6/5.4/7.2 ms with 6/8 slice partial Fourier; volumetric navigators, vNav, for prospective head-motion correction with, FOV = 256x240 (93.8%) mm.

### Data analysis

Anatomical and functional images were preprocessed using fMRIprep (Esteban et al., 2019; see Supplementary Material for full details).

#### GLM

Surface based, fsaverage space cleaned timeseries outputs from fmriprep were read into MATLAB 2019a for further analysis (MathWorks Inc., Natick, USA). GLM single-trial beta estimates were computed using the GLMsingle toolbox (Prince et al., 2022; Kay et al., 2013; Rokem & Kay, 2020) and used as a basis for both MVPA and univariate analysis. The GLMsingle algorithm provides several advantages over a typical GLM. The first is the use of a library of HRFs, whereby the best-fitting HRF from the library is chosen for each voxel and provides well-regularized HRF estimates. The second uses the repeated runs of the design to generate data-derived nuisance regressors to remove noise from beta estimates. The third is an application of ridge regression as a method for dampening the noise inflation caused by correlated single-trial GLM predictors. To determine the optimal level of regularization for each voxel fractional ridge regression is used (Rokem & Kay, 2020).

The extracted betas for each voxel represented estimates of the trial-wise BOLD response amplitudes to each stimulus trial (4 seconds of sound presentation), and these were relative to the BOLD signal observed during the absence of a stimulus (silence). A normalized version of the betas was used for carrying out MVPA analysis. Instead, prior to running the univariate analyses, we spatially smoothed the betas from first-level models for each participant. Results of the univariate analyses are reported in Supplementary Material.

#### ROI definition

Predefined Regions of Interest (ROIs) consisted of V1, precuneus (PC) the Early Auditory Cortex (EAC). All regions were defined using the Human Connectome Project-MultiModal Parcellation atlas (HCP-MMP, Glasser et al., 2016). ROIs were combined across both hemispheres. The precuneus ROI was defined as the Precuneus Visual Area of the HCP-MMP atlas, which corresponds to the anterior/ventral precuneus (Jitsuishi & Yamaguchi, 2023). This specific region has been implicated in recovery of consciousness (Wu et al., 2022), imagery (Fulford et al., 2018; Jian et al., 2015), memory (Richter et al., 2016; Mazzoni et al., 2019) and attention (Natale et al., 2006) and it is characterized by direct connections with the visual cortex (Rolls et al., 2022). The Early Auditory Cortex (EAC) was composed of the primary auditory cortex (A1) and the lateral and medial belts (MBelt, LBelt), following the guidelines of HCP-MMP atlas (Glasser et al., 2016).

#### ROI-based MVPA

MVPA was performed with the CoSMo MVPA Toolbox for MATLAB (available at http://www.cosmomvpa.org, Oosterhof et al., 2016). For each participant, single trial normalized beta weights for all tasks were extracted from all vertices within an ROI. A linear support vector machine classification algorithm (LIBSVM toolbox, Chang & Lin, 2011) was used to decode activity patterns from individual runs. A ‘leave-one-run-out’ cross-validation approach was used to estimate the accuracy of the classifier. Specifically, for each task (passive listening and voluntary imagery) the classifier was trained on 4 runs to distinguish between the six categories of sounds (one-versus-one classification, e.g., dog vs seagulls vs cat vs helicopter vs fireplace vs traffic) and tested on the remaining 5th run and the results were averaged. As recommended by Stelzer, Chen & Turner (2013), we assessed the statistical significance of classifier performance by performing a permutation analysis on all classifications and for each ROI (V1, PC and EAC as control region). This analysis provides a more robust and sensitive test of statistical significance compared to a one-sample t-test against the theoretical chance level. Specifically, we trained and tested the classifier across 1000 permutations with randomised labels in each subject and each ROI (V1, PC, EAC). On the group level, following the same approach of Vetter et al. (2014, 2020), we computed p-values from the mean randomisation distribution and the mean real label performance. Specifically, p-values were calculated as the probability of obtaining a value as large as the real label performance in the randomisation distribution, yielding a minimum p-value of 0.001 (Vetter et al., 2014).

#### Statistical analyses

Further statistical analyses were carried out in R (R Core Team, 2022). We conducted a linear mixed effect model analysis (LMM, West et al., 2006) to test for differences in subjective vividness ratings or decoding accuracy in our ROIs across groups and tasks. The LMM analysis was performed using the lmer() function of the lme4 library (Bates et al., 2014). Significance was calculated using the lmerTest package (Kuznetsova et al., 2017), which applies Satterthwaite’s method to estimate degrees of freedom and generate p-values for mixed models. We used the emmeans package (Lenth, 2020) to compute post-hoc comparisons with Tukey-adjusted p-values. Our models included decoding accuracy in one of our ROIs or vividness ratings as the outcome variable, while task condition (passive listening task, reflecting spontaneous imagery, and voluntary imagery task, reflecting voluntary imagery) and group (aphantasics, visualizers), as well as their interaction, were included as fixed effects. Finally, participant identifiers were included as random effects. We also carried out a correlational analysis using Pearson’s correlation coefficient (r) to examine whether there was a relationship between vividness ratings and decoding accuracy in V1 for voluntary imagery and passive listening tasks in visualizers.

## Footnotes

1 Zeman et al. (2015) reported that 10/11 aphantasic participants with a complete absence of subjective voluntary imagery described experiencing visual imagery in dreams ‘and/or’ ‘brief flashes of imagery’ during wakefulness. However, while Zeman et al. interpreted the latter as reflecting involuntary imagery, we point out that “brief flashes” still equate to the definition of aphantasia (i.e., imagery is fully absent or vague/dim/fleeting). The key difference may therefore be, very simply, that their fleeting imagery didn’t happen to arise during the 3 minutes they spent completing the voluntary imagery questionnaire (because this was a tiny portion of their waking life).

