## Supplementary Material for "Sensory representations in primary visual cortex are not sufficient for subjective imagery"

### Image pre-processing

Results included in this manuscript come from preprocessing performed using fMRIPrep 21.0.0 (Esteban, et al., 2018; Esteban, Blair, et al. 2018); RRID:SCR_016216), which is based on Nipype 1.6.1 (K. Gorgolewski et al. 2011; K. J. Gorgolewski et al., 2018); RRID:SCR_002502).

*Preprocessing of B0 inhomogeneity mappings*

A total of 1 fieldmaps were found available within the input BIDS structure for this particular subject. A B0-nonuniformity map (or fieldmap) was estimated based on two (or more) echo-planar imaging (EPI) references with topup (Andersson, Skare, and Ashburner, 2003); FSL 6.0.5.1:57b01774).

*Anatomical data preprocessing*

A total of 1 T1-weighted (T1w) images were found within the input BIDS dataset. The T1-weighted (T1w) image was corrected for intensity non-uniformity (INU) with N4BiasFieldCorrection (Tustison et al. 2010), distributed with ANTs 2.3.3 (Avants et al. 2008, RRID:SCR_004757), and used as T1w-reference throughout the workflow. The T1w-reference was then skull-stripped with a Nipype implementation of the antsBrainExtraction.sh workflow (from ANTs), using OASIS30ANTs as target template. Brain tissue segmentation of cerebrospinal fluid (CSF), white-matter (WM) and gray-matter (GM) was performed on the brain-extracted T1w using fast (FSL 6.0.5.1:57b01774, RRID:SCR_002823, Zhang, Brady, and Smith 2001). Brain surfaces were reconstructed using recon-all (FreeSurfer 6.0.1, RRID:SCR_001847, Dale, Fischl, and Sereno 1999), and the brain mask estimated previously was refined with a custom variation of the method to reconcile ANTs-derived and FreeSurfer-derived segmentations of the cortical gray-matter of Mindboggle (RRID:SCR_002438, Klein et al. 2017). Volume-based spatial normalization to two standard spaces (MNI152NLin6Asym, MNI152NLin2009cAsym) was performed through nonlinear registration with antsRegistration (ANTs 2.3.3), using brain-extracted versions of both T1w reference and the T1w template. The following templates were selected for spatial normalization: FSL's MNI ICBM 152 non-linear 6th Generation Asymmetric Average Brain Stereotaxic Registration Model [Evans et al. (2012), RRID:SCR_002823; TemplateFlow ID: MNI152NLin6Asym], ICBM 152 Nonlinear Asymmetrical template version 2009c [Fonov et al. (2009), RRID:SCR_008796; TemplateFlow ID: MNI152NLin2009cAsym].

*Functional data preprocessing*

For each of the 5 BOLD runs found per subject (across all tasks and sessions), the following preprocessing was performed. First, a reference volume and its skull-stripped version were generated by aligning and averaging 1 single-band references (SBRefs). Head-motion parameters with respect to the BOLD reference (transformation matrices, and six corresponding rotation and translation parameters) are estimated before any spatiotemporal filtering using mcflirt (FSL 6.0.5.1:57b01774, Jenkinson et al. 2002). The estimated fieldmap was then aligned with rigid-registration to the target EPI (echo-planar imaging) reference run. The field coefficients were mapped on to the reference EPI using the transform. BOLD runs were slice-time corrected to 0.719s (0.5 of slice acquisition range 0s-1.44s) using 3dTshift from AFNI (Cox and Hyde 1997, RRID:SCR_005927). The BOLD reference was then co-registered to the T1w reference using bbregister (FreeSurfer) which implements boundary-based registration (Greve and Fischl 2009). Co-registration was configured with six degrees of freedom. First, a reference volume and its skull-stripped version were generated using a custom methodology of fMRIPrep. Several confounding time-series were calculated based on the preprocessed BOLD: framewise displacement (FD), DVARS and three region-wise global signals. FD was computed using two formulations following Power (absolute sum of relative motions, Power et al. (2014)) and Jenkinson (relative root mean square displacement between affines, Jenkinson et al. (2002)). FD and DVARS are calculated for each functional run, both using their implementations in Nipype (following the definitions by Power et al. 2014). The three global signals are extracted within the CSF, the WM, and the whole-brain masks. Additionally, a set of physiological regressors were extracted to allow for component-based noise correction (CompCor, Behzadi et al. 2007). Principal components are estimated after high-pass filtering the preprocessed BOLD time-series (using a discrete cosine filter with 128s cut-off) for the two CompCor variants: temporal (tCompCor) and anatomical (aCompCor). tCompCor components are then calculated from the top 2% variable voxels within the brain mask. For aCompCor, three probabilistic masks (CSF, WM and combined CSF+WM) are generated in anatomical space. The implementation differs from that of Behzadi et al. in that instead of eroding the masks by 2 pixels on BOLD space, the aCompCor masks are subtracted a mask of pixels that likely contain a volume fraction of GM. This mask is obtained by dilating a GM mask extracted from the FreeSurfer's aseg segmentation, and it ensures components are not extracted from voxels containing a minimal fraction of GM.

Finally, these masks are resampled into BOLD space and binarized by thresholding at 0.99 (as in the original implementation). Components are also calculated separately within the WM and CSF masks. For each CompCor decomposition, the k components with the largest singular values are retained, such that the retained components' time series are sufficient to explain 50 percent of variance across the nuisance mask (CSF, WM, combined, or temporal). The remaining components are dropped from consideration. The head-motion estimates calculated in the correction step were also placed within the corresponding confounds file. The confound time series derived from head motion estimates and global signals were expanded with the inclusion of temporal derivatives and quadratic terms for each (Satterthwaite et al. 2013). Frames that exceeded a threshold of 0.5 mm FD or 1.5 standardised DVARS were annotated as motion outliers. The BOLD time-series were resampled into several standard spaces, correspondingly generating the following spatially-normalized, preprocessed BOLD runs: MNI152NLin6Asym, MNI152NLin2009cAsym. First, a reference volume and its skull-stripped version were generated using a custom methodology of fMRIPrep. The BOLD time-series were resampled onto the following surfaces (FreeSurfer reconstruction nomenclature): fsaverage. Automatic removal of motion artifacts using independent component analysis (ICA-AROMA, Pruim et al. 2015) was performed on the preprocessed BOLD on MNI space time-series after removal of non-steady state volumes and spatial smoothing with an isotropic, Gaussian kernel of 6mm FWHM (full-width half-maximum). Corresponding "non-aggresively" denoised runs were produced after such smoothing. Additionally, the "aggressive" noise-regressors were collected and placed in the corresponding confounds file. All resamplings can be performed with a single interpolation step by composing all the pertinent transformations (i.e. head-motion transform matrices, susceptibility distortion correction when available, and co-registrations to anatomical and output spaces). Gridded (volumetric) resamplings were performed using antsApplyTransforms (ANTs), configured with Lanczos interpolation to minimize the smoothing effects of other kernels (Lanczos 1964). Non-gridded (surface) resamplings were performed using mri_vol2surf (FreeSurfer).

*Functional data preprocessing*

For each of the 5 BOLD runs found per subject (across all tasks and sessions), the following preprocessing was performed. First, a reference volume and its skull-stripped version were generated by aligning and averaging 1 single-band references (SBRefs). Head-motion parameters with respect to the BOLD reference (transformation matrices, and six corresponding rotation and translation parameters) are estimated before any spatiotemporal filtering using mcflirt (FSL 6.0.5.1:57b01774, Jenkinson et al. 2002). The estimated fieldmap was then aligned with rigid-registration to the target EPI (echo-planar imaging) reference run. The field coefficients were mapped on to the reference EPI using the transform. BOLD runs were slice-time corrected to 0.718s (0.5 of slice acquisition range 0s-1.44s) using 3dTshift from AFNI (Cox and Hyde 1997, RRID:SCR_005927). The BOLD reference was then co-registered to the T1w reference using bbregister (FreeSurfer) which implements boundary-based registration (Greve and Fischl 2009). Co-registration was configured with six degrees of freedom. First, a reference volume and its skull-stripped version were generated using a custom methodology of fMRIPrep. Several confounding time-series were calculated based on the preprocessed BOLD: framewise displacement (FD), DVARS and three region-wise global signals. FD was computed using two formulations following Power (absolute sum of relative motions, Power et al. (2014)) and Jenkinson (relative root mean square displacement between affines, Jenkinson et al. (2002)). FD and DVARS are calculated for each functional run, both using their implementations in Nipype (following the definitions by Power et al. 2014). The three global signals are extracted within the CSF, the WM, and the whole-brain masks. Additionally, a set of physiological regressors were extracted to allow for component-based noise correction (CompCor, Behzadi et al. 2007). Principal components are estimated after high-pass filtering the preprocessed BOLD time-series (using a discrete cosine filter with 128s cut-off) for the two CompCor variants: temporal (tCompCor) and anatomical (aCompCor). tCompCor components are then calculated from the top 2% variable voxels within the brain mask. For aCompCor, three probabilistic masks (CSF, WM and combined CSF+WM) are generated in anatomical space. The implementation differs from that of Behzadi et al. in that instead of eroding the masks by 2 pixels on BOLD space, the aCompCor masks are subtracted a mask of pixels that likely contain a volume fraction of GM. This mask is obtained by dilating a GM mask extracted from the FreeSurfer's aseg segmentation, and it ensures components are not extracted from voxels containing a minimal fraction of GM. Finally, these masks are resampled into BOLD space and binarized by thresholding at 0.99 (as in the original implementation). Components are also calculated separately within the WM and CSF masks. For each CompCor decomposition, the k components with the largest singular values are retained, such that the retained components' time series are sufficient to explain 50 percent of variance across the nuisance mask (CSF, WM, combined, or temporal). The remaining components are dropped from consideration. The head-motion estimates calculated in the correction step were also placed within the corresponding confounds file. The confound time series derived from head motion estimates and global signals were expanded with the inclusion of temporal derivatives and quadratic terms for each (Satterthwaite et al. 2013). Frames that exceeded a threshold of 0.5 mm FD or 1.5 standardised DVARS were annotated as motion outliers. The BOLD time-series were resampled into several standard spaces, correspondingly generating the following spatially-normalized, preprocessed BOLD runs: MNI152NLin6Asym, MNI152NLin2009cAsym. First, a reference volume and its skull-stripped version were generated using a custom methodology of fMRIPrep. The BOLD time-series were resampled onto the following surfaces (FreeSurfer reconstruction nomenclature): fsaverage. Automatic removal of motion artifacts using independent component analysis (ICA-AROMA, Pruim et al. 2015) was performed on the preprocessed BOLD on MNI space time-series after removal of non-steady state volumes and spatial smoothing with an isotropic, Gaussian kernel of 6mm FWHM (full-width half-maximum). Corresponding "non-aggresively" denoised runs were produced after such smoothing. Additionally, the "aggressive" noise-regressors were collected and placed in the corresponding confounds file. All resamplings can be performed with a single interpolation step by composing all the pertinent transformations (i.e. head-motion transform matrices, susceptibility distortion correction when available, and co-registrations to anatomical and output spaces). Gridded (volumetric) resamplings were performed using antsApplyTransforms (ANTs), configured with Lanczos interpolation to minimize the smoothing effects of other kernels (Lanczos 1964). Non-gridded (surface) resamplings were performed using mri_vol2surf (FreeSurfer).

Many internal operations of fMRIPrep use Nilearn 0.8.1 (Abraham et al. 2014, RRID:SCR_001362), mostly within the functional processing workflow. For more details of the pipeline, see the section corresponding to workflows in fMRIPrep's documentation.

*Copyright Waiver*

The above boilerplate text was automatically generated by fMRIPrep with the express intention that users should copy and paste this text into their manuscripts unchanged. It is released under the CC0 license.

**Relationship between Vividness of Visual Imagery Questionnaire (VVIQ; Marks, 1973) or Spontaneous Use of Imagery Scale (SUIS; Reisberg et al., 2003) and decoding accuracy of sounds in primary visual cortex (V1).**

In visualizers, decoding accuracy in V1 during voluntary imagery correlated with individual VVIQ scores (Figure S1A), $r(24) = .42$, $p = .03$, in line with the results observed for the ratings of vividness collected during scanning. No significant correlation was observed between VVIQ and decoding accuracy in V1 for passive listening, $r(24) = .17$, $p = .41$, Figure S1B. Within the same group, we also carried out an exploratory analysis to test whether participants' scores on the SUIS, which reflects an index of frequency of mental imagery use in daily life, were related to decoding accuracy of sounds content in V1. We found no significant relationship between SUIS and decoding accuracy in V1 for either passive listening ($r = .32$, $p = .11$, Figure S1C) and voluntary ($r = .09$, $p = .66$, Figure S1D) imagery tasks.

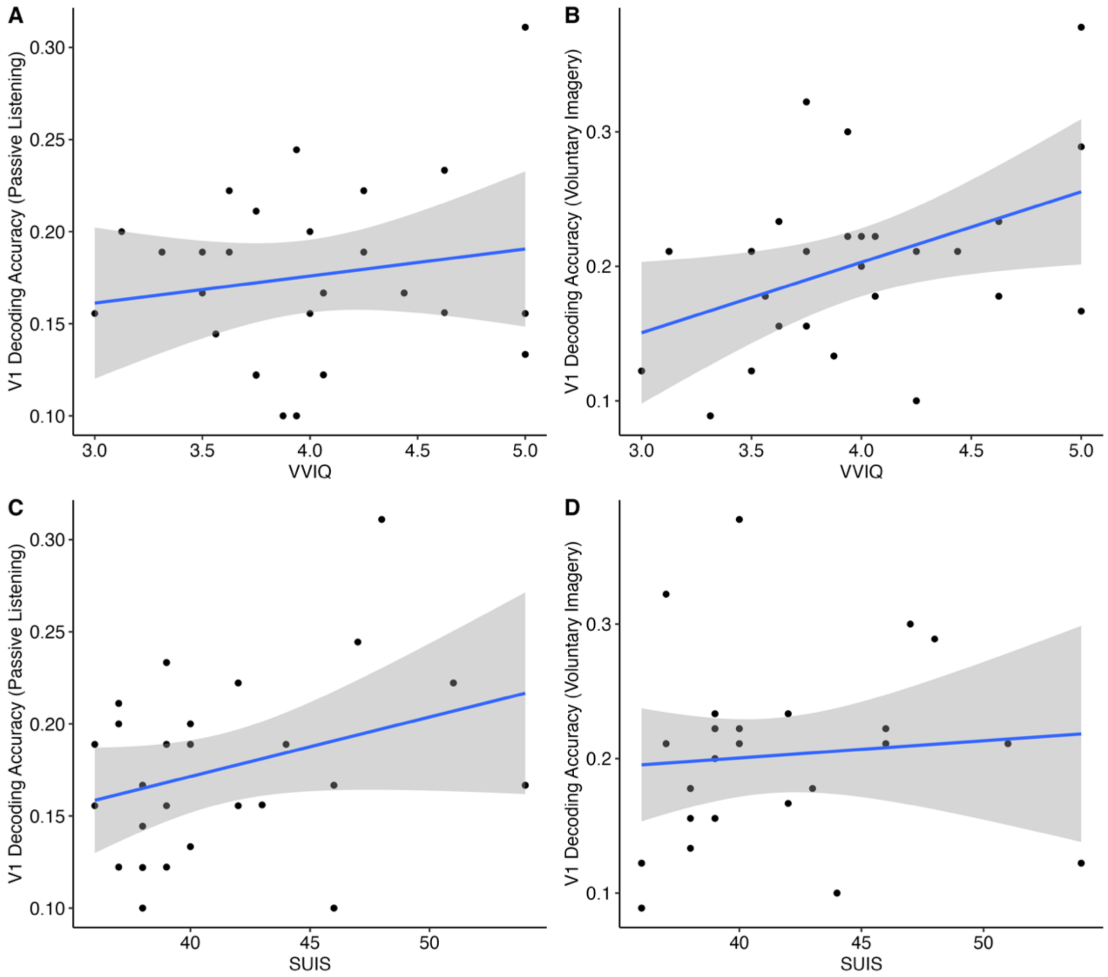


**Figure S1.** Relationship between VVIQ (A, B) or SUIS (C, D) and decoding accuracy in V1 during passive listening task (A, C) or voluntary imagery task (B, D) in the visualizer group. Shaded interval shows 95% CI.

**Univariate ROI results**

We compared activity induced by listening to sounds and by blank periods of silence (except for MRI scanner noise) during both passive listening and voluntary imagery task. In particular, we averaged betas extracted from our ROIs (V1, PC, EAC) for the contrast of all sounds vs blank periods (no sound). First, we compared differences in V1 activation for this contrast between tasks and groups and found no main effect of task, $F(1, 48) = 2.68$, $p = .11$, $\eta^2_p = 0.05$, group, $F(1, 48) = 1.00$, $p = .32$, $\eta^2_p = 0.02$, or interaction between the two, $F(1, 48) = 1.33$, $p = .25$, $\eta^2_p = 0.03$. As it can be seen in Figure S2A, similarly to Vetter et al. (2014), V1 was characterized by a slight deactivation in both groups and tasks.

In precuneus ROI, no main effect of group was observed, $F(1, 48) = 0.04$, $p = .84$. However, we observed a significant main effect of task, $F(1, 48) = 14.20$, $p < .001$, $\eta^2_p = 0.23$, with PC being more deactivated during voluntary imagery ($M = -0.05$, $SD = 0.08$) than during passive listening ($M = -0.02$, $SD = 0.07$), see Figure S2B. No significant interaction was observed between task and group, $F(1, 48) = 2.30$, $p = .14$, $\eta^2_p = 0.05$.

Regarding EAC, as shown in Figure S2C, we found a main effect of task, $F(1, 48) = 8.51$, $p = .005$, $\eta^2_p = 0.15$, reflecting higher activation during the voluntary imagery task ($M = 1.08$, SD = 0.5) than during passive listening ($M = 0.98$, $SD = 0.47$), in line with multivariate results, but no main effect of group, $F(1, 48) = 0.003$, $p = .95$, nor interaction between the two, $F(1, 48) = 1.00$, $p = .32$, $\eta^2_p = 0.02$.

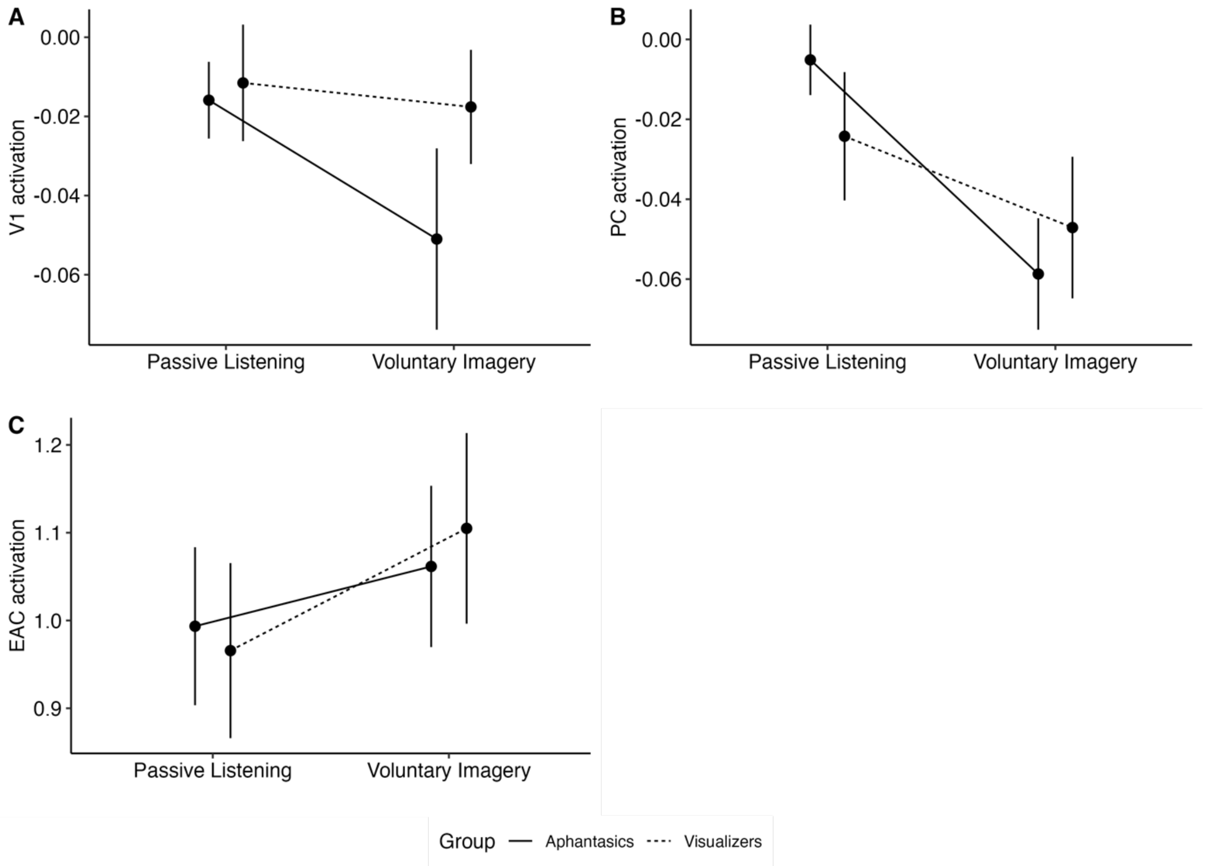


**Figure S2.** ROI activation. **A.** Primary Visual Cortex (V1), **B.** Precuneus (PC), **C.** Early Auditory Cortex (EAC) activation by group (aphantasic, control) and task condition (passive listening, voluntary imagery). Error bars show SEM.

### Pilot Studies for evocative sounds selection

We conducted three online pilot studies to select the series of natural sounds that would be used to evoke visual mental imagery in the main fMRI study. The purpose of this pilot work was to select sound stimuli that would be effective in inducing spontaneous visual imagery during the first part of the scanning session (passive listening), without explicitly instructing or biasing our participant in any way to conjure visual imagery (as this would inevitably affect the nature of the imagery generated and reduce its “spontaneity”). Across the three pilots, we gathered ratings related to spontaneous and voluntary imagery vividness and tested different categorizations of sounds and/or their relationship with VVIQ scores.

***Pilot 1***

The first online pilot served as initial pre-screening of sound stimuli from a pool of 101 natural sounds, selected from pre-existing databases (the IADS-E, Yang et al., 2018, and IADS-2, Bradley & Lang, 2007a) and online sources (e.g., https://freesound.org/). 66 participants (age range 18-41, mean age 21.3 ± 3.66) listened to all the sounds and provided the following ratings for each sound: the duration of the visual imagery they experienced during listening (from 1, “never”, to 4, “all the time”), vividness of visual imagery evoked by the sound (from 1, “no image at all”, to 5, “perfectly clear and vivid”), and identification of the sound as being produced by a living or non-living entity. In the instructions, we explicitly indicated that we were interested in assessing spontaneous imagery evoked by natural sounds. Importantly, we told our participants that they did not need to make an effort or actively try to engage in visual imagery while listening to the sounds. Instead, participants were instructed to just listen carefully to each sound and report whether any image associated with the sounds’ content spontaneously emerged in their mind.

We first excluded all the sounds that were assigned to the corresponding category (living, non-living) by less than 85% of participants. Among the remaining 85 sounds, we selected 8 “high vividness” sounds (4 living, 4 non-living) and 8 “low vividness” sounds (4 living, 4 non-living). The selected sound categories were: “cat”, “airplane”, “crickets”, “dog”, “ambulance”, “drilling”, “duck”, “fire”, “frog”, “helicopter”, “seagulls”, “traffic”, “turkey”, “washing machine”, “wind”, “dolphin”.

***Pilot 2***

Our second pilot examined whether the categorization in “low” and “high” vividness of the sounds selected in Pilot 1 could generalize to a new sample of participants (at the time of piloting, we were considering the possibility of a within-subject manipulation of imageability). Moreover, here we tested the relationship between sound-specific vividness ratings of spontaneous and voluntary imagery with participants’ VVIQ score. The ratings were gathered within a paradigm similar to the one we planned to use in the fMRI study. Specifically, the spontaneous imagery rating consisted of a single rating of vividness (from 1, “no image at all”, to 5, “perfectly clear and vivid”) collected after a first listening of all the sounds. This was followed by a second listening of all the sounds during which participants were requested to voluntarily generate visual mental images of what was producing the sound and rate their vividness. 75 participants completed the task (mean age 20.01 ± 1.47, 66 females, 8 males, 1 other/prefer not to say), 38 in the high and 37 in the low imageable sounds condition. In addition to the selected 8 high vividness (living, non-living) and 8 low vividness sound categories, we also included for each category two additional exemplars (e.g., another exemplar of a dog barking). This was done to have three exemplars for each sound category. We selected the two different exemplars for each sound category from the same sources as Pilot 1, for a total of 48 sound exemplars being tested.

*Results.* First, we checked whether our two groups of participants significantly differed in their imagery abilities as indexed by the VVIQ. No difference was observed, $t(72.19) = -0.93$, $p = .35$. We then tested for differences in ratings of vividness between the high and low imageable sounds group. Contrary to our expectations, there was no significant difference between the two groups for spontaneous, $t(72.79) = 0.37$, $p = .35$ (Figure S3A), and voluntary imagery vividness ratings, $t(72.69) = 0.96$, $p = .16$ (Figure S3B). On the other hand, we found that across participants, VVIQ positively correlated with both spontaneous, $r(73) = .49$, $p < .001$ (Figure S3C), and voluntary imagery vividness, $r(73) = .71$, $p < .001$ (Figure S3D), suggesting that the clarity of sound-evoked imagery was driven by individual differences in subjective imagery abilities, rather than our predefined categorization in high and low imageable sounds.

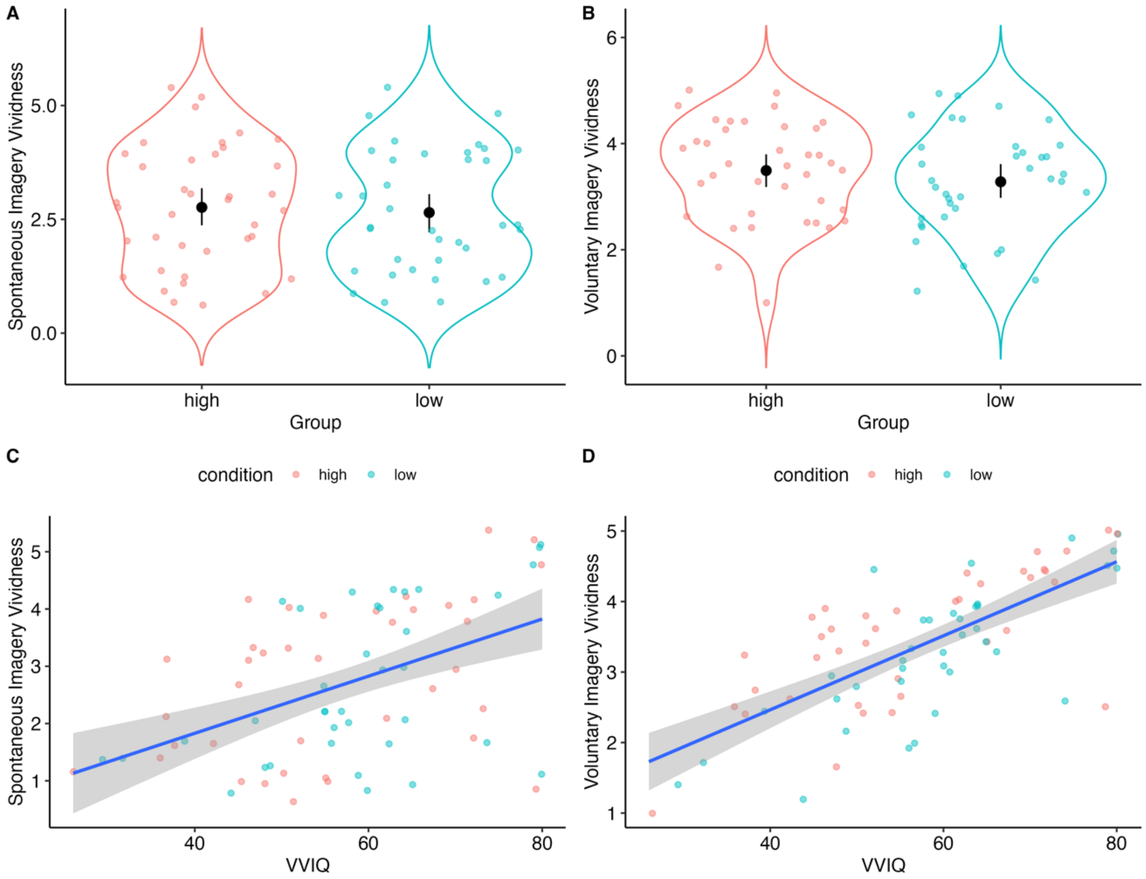


**Figure S3. A.** Spontaneous (single rating) and **B.** voluntary imagery (mean rating across sounds per participant) ratings as a function of group. Dots represent individual participants. Error bar shows 95 % CI. **C.** Spontaneous and **D.** voluntary subjective imagery ratings as a function of VVIQ (points represent single subjects, filled with colors of the respective group). Shaded interval shows 95% CI.

### *Pilot 3*

Results of Pilot 2 suggested that the imageability of sounds was highly dependent on self-reported imagery abilities. In contrast, no effect of categorization into low and high vividness sounds was observed between participants. For our main data collection (fMRI study) we therefore decided to drop categorization in high and low imageable sounds and rather rely on self-reported imagery abilities as indexed by the VVIQ. In a third pilot, we conducted a final selection of sound stimuli by testing their vividness again as well as considering whether participants could correctly indicate the name corresponding to the entity or object producing the sound. Specifically, we tested the similarity index between the name that was originally assigned to each sound category (e.g., dog, duck, seagulls) and the name that would be indicated by participants for each sound exemplar during the experimental session. 35 participants (mean age

19.83 ± 1.56, 29 females, 6 males) took part in Pilot 3. Participants were asked to listen carefully to the sounds and, at the end of each sound presentation, answer the following questions: 1) "How vivid was the visual imagery that you experienced while listening to the sound?", on a Likert scale from 1 (no image at all) to 5 (perfectly clear and vivid); 2) "Can you name what the sound represents (what is producing the sound or what scenario do you associate with the sound)?". Participants could answer Question 2 by typing the chosen name inside a box (free text) and were recommended to use a single name to indicate the entity producing the sound.

*Results.* We calculated a similarity score between the word indicating the entity producing the sound provided by participants with the word that we used to define our sound categories. To do so, we used the package stringdist (Van der Loo, 2014) implemented in R, which allows the computation of metrics indexing approximate text matching between different string vectors. We specifically employed the Jaro-Winkler Similarity, a metric recommended for matching short strings, mainly names or entity names (Jaro, 1989), that ranges from 0 to 1. A similarity score was calculated for each of the 48 sounds exemplars within each participant, by comparing the name provided by the participant with the name that we originally assigned to the sound (e.g., dog). We then averaged mean similarity and vividness across participants for each sound exemplar. Our initial selection comprised 4 living (cat, dog, seagulls, duck) and 4 non-living (traffic, wind, fire, helicopter) sound categories with the highest similarity and vividness scores. After testing one pilot participant in the scanner, we removed two sound categories (duck, wind) as their respective sound exemplars could not be clearly heard by the pilot participant while the scanner was functioning. The final sound categories we selected were cat, dog, seagulls, fire, traffic, and helicopter. Table S1 reports vividness and mean similarity for the final selection of sound categories with respective exemplars.

**Table S1**

*Mean ratings of similarity and vividness of the final set of sounds selected as main stimuli in our main data collection. 'Vividness' consists of a rating on a Likert scale ranging between ("no image at all") to 5 ("perfectly clear and vivid"). 'Similarity' is a score calculated by the overlap between the sound as named by the participant and the designated category name (e.g., in the case of the cat sounds, a response 'cat' would have a similarity score of 1).*

| Category | Exemplar | Similarity | Vividness |
|---|---|---|---|
| cat | cat1 | 0.93 | 3.80 |
| cat | cat2 | 0.96 | 3.86 |
| cat | cat3 | 0.94 | 3.74 |
| dog | dog1 | 0.94 | 4.03 |
| dog | dog2 | 0.94 | 4.11 |
| dog | dog3 | 0.91 | 4.03 |
| fire | fire1 | 0.91 | 3.23 |
| fire | fire2 | 0.87 | 3.54 |
| fire | fire3 | 0.95 | 3.66 |
| helicopter | helicopter1 | 0.95 | 3.71 |
| helicopter | helicopter2 | 0.80 | 3.49 |
| helicopter | helicopter3 | 0.93 | 3.51 |
| seagulls | seagulls1 | 0.94 | 4.29 |
| seagulls | seagulls2 | 0.92 | 4.31 |
| seagulls | seagulls3 | 0.94 | 4.34 |
| traffic | traffic1 | 0.82 | 4.06 |
| traffic | traffic2 | 0.87 | 3.97 |
| traffic | traffic3 | 0.88 | 3.94 |
